# One critic, two actors: Evidence for covert learning in the basal ganglia

**DOI:** 10.1101/060236

**Authors:** Meropi Topalidou, Daisuke Kase, Thomas Boraud, Nicolas P. Rougier

## Abstract

This paper introduces a new hypothesis concerning the dissociated role of the basal ganglia in the selection and the evaluation of action that has been formulated using a theoretical model and confirmed experimentally in monkeys. To do so, prior to learning, we inactivated the internal part of the Globus Pallidus (GPi, the main output structure of the BG) with injections of muscimol and we tested monkeys on a variant of a two-armed bandit task where two stimuli are associated with two distinct reward probabilities (0.25 and 0.75 respectively). Unsurprisingly, performance in such condition are at the chance level because the output of basal ganglia is suppressed and they cannot influence behaviour. However, the theoretical model predicts that in the meantime, values of the stimuli are nonetheless covertly evaluated and learned. This has been tested and confirmed on the next day, when muscimol has been replaced by a saline solution: monkeys instantly showed significantly improved performances (above chance level), hence demonstrating they have covertly learned the relative value of the two stimuli. This tends to suggest a competition takes place in the Cortex-BG loop between two actors, one of whom being sensitive to criticism and the other not. Ultimately, the actual choice is valuated, independently of the origin of the decision.

## Introduction

The now classical actor-critic model of decision making elaborated in the 1980s posits that there are two separate components in order to explicitly represent the policy independently of the value function. The actor is in charge of choosing an action in a given state (policy) while the critic is in charge of evaluating (criticizing) the current state (value function). This classical view has been used extensively for modelling the basal ganglia (Suri and Schultz, 1999; Suri, 2002; Doya, 2007; Glimcher, 2011; Doll et al., 2012) even though the precise anatomical mapping of these two components is still subject to debate and may diverge from one model to the other (Redgrave et al., 2008; Niv and Langdon, 2016). However, all these models share the implicit assumption that the actor and the critic are acting in concert, i.e. the actor determines the policy exclusively from the values estimated by the critic, as in Q-Learning or SARSA. Interestingly enough, (Sutton and Barto, 1998) noted in their seminal work that *one could imagine intermediate architectures in which both an action-value function and an independent policy would be learned*. One legitimate question is thus to wonder whether the role of the BG critic is restricted to the exclusive evaluation of the BG actor or if this role might be considered to be more general. In other words, can the BG critic role extend beyond the basal ganglia and evaluate any actor, independently on its origin?

We support this latter hypothesis based on a decision-making model that is grounded on anatomical and physiological data and that identify the cortex-basal ganglia (CBG) loop as the actor. The critic — of which the Substantia Nigra pars compacta (SNc) and the Ventral Tegmental Area (VTA) are essential components — interacts through dopamine projections to the striatum (Leblois et al., 2006). Decision is generated by symmetry breaking mechanism that emerges from competitions processes between positives and negatives feedback loop encompassing the full CBG network (Guthrie et al., 2013). This model captured faithfully behavioural, electrophysiological and pharmacological data we obtained in primates using implicit variant of two-armed bandit tasks — that assessed both learning and decision making — but was less consistent with the explicit version (i.e. when values are known from the beginning of the task) that focus on the decision process only. We therefore upgraded this model by adding a cortical module that is granted with a competition mechanism and Hebbian learning (Doya, 2000). This improved version of the model predicts that the whole CBG loop is actually necessary for the implicit version of the task, however, when the basal ganglia feedback to cortex is disconnected, the system is still able to choose in the explicit version of the task. Our experimental data fully confirmed this prediction (Piron et al., 2016) and allowed to solve an old conundrum concerning the pathophysiology of the BG which was that lesion or jamming of the output of the BG improve parkinsonian patient motor symptoms while it affects marginally their cognitive and psychomotor performances.

An interesting prediction of this generalized actor-critic architecture is that the valuation of options and the behavioural outcome are segregated. In the computational model, it implies that if we block the output of the basal ganglia in a two-armed bandit task before learning, and because reinforcement learning occurred at the striatal level under dopaminergic control, this should induce covert learning when the model chooses randomly. The goal of this study is thus twofold: i) to present a comprehensive description of the model in order to provide the framework for an experimental paradigm that allow to objectivize covert learning and ii) to test this prediction in monkeys.

## Material and methods

### Behavioral experiments

Experimental procedures were performed in accordance with the Council Directive of 20 October 2010 (2010/63/UE) of the European Community. This project was approved by the French Ethic Comity for Animal Experimentation (#50120111-A). Data were obtained from two female macaque monkeys that were previously used in a related set of experiments. All the details concerning animal care, experimental setup, surgical procedure, bilateral inactivation of the GPi and histology can be found in (Piron et al., 2016). Raw data is available from (Kase and Boraud, 2017)

### Computational modeling

Code and statistical analysis are available from (Rougier and Topalidou, 2017).

#### Architecture

The model is an extension of previously published models (Leblois et al., 2006; Guthrie et al., 2013). The model by (Leblois et al., 2006) introduced an action selection mechanism which derives from the competition between a positive feedback through the direct pathway and a negative feedback through the hyper-direct pathway in the cortico-basal-thalamic loop. The model has been extended in (Guthrie et al., 2013) in order to explore the parallel organization of circuits in the BG. This model includes all the major nuclei of the basal ganglia (but GPe) and is organized along three segregated loops (motor, associative and cognitive) that spread over the cortex, the basal ganglia and the thalamus (Alexander et al., 1986; Albin et al., 1989; Alexander and Crutcher, 1990; Parent and Hazrati, 1995). It incorporates a two-level decision making with a cognitive level selection (lateral prefrontal cortex, LPFC) based on cue shape and a motor level selection (supplementary motor area, SMA, and primary motor cortex, PMC) based on cue position (see figure 2). In this latter model (Rougier and Topalidou, 2017), the cortex was mostly an input/output structure under the direct influence of both the task input and the thalamic output resulting from the basal ganglia computations. Consequently, this cortex could not take a decision of its own. In the present work, and to cope with our main hypothesis, we added a lateral competition mechanism in all three cortices (motor, cognitive, associative) based on short range excitation and long range inhibition and connections (Coombes et al., 2014) between the associative cortex and cognitive (resp. motor ones) to allow for the cross talking of these structures. This competition results in the capacity for the cortex to make a decision, although with a slower dynamic when compared to the BG.

**Figure 1.**
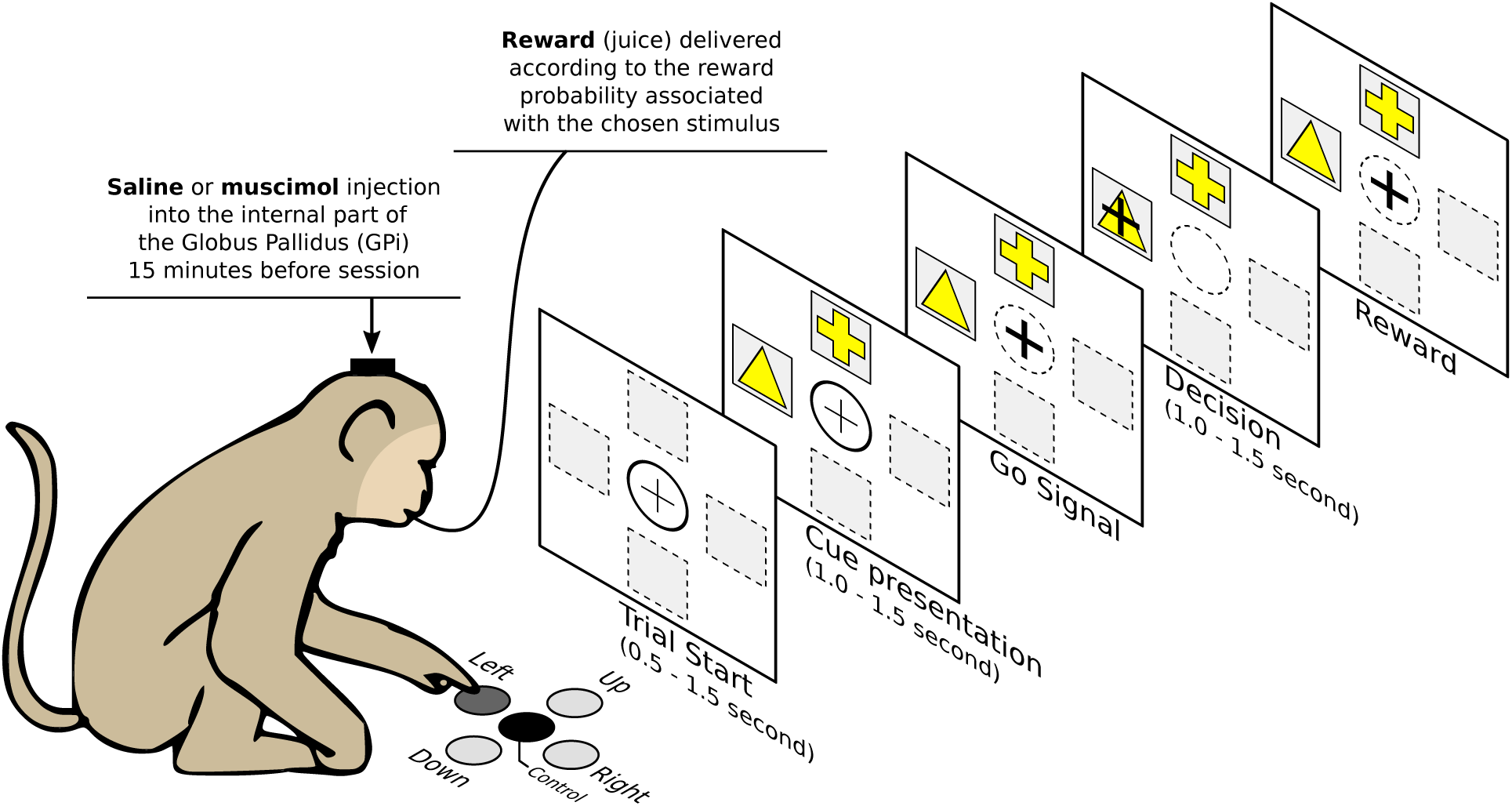
A trial is made of the simultaneous presentation of two cues at two random positions associated with a fixed reward probability. The monkey has to choose a stimulus at the go signal and maintain this choice for one second. Reward is delivered according to the reward probability associated with the chosen stimulus. For all the experiments in this study, we used fixed reward probability set respectively to 0.75 and 0.25. Figure from (Rougier, 2017), CC-BY.

**Figure 2.**
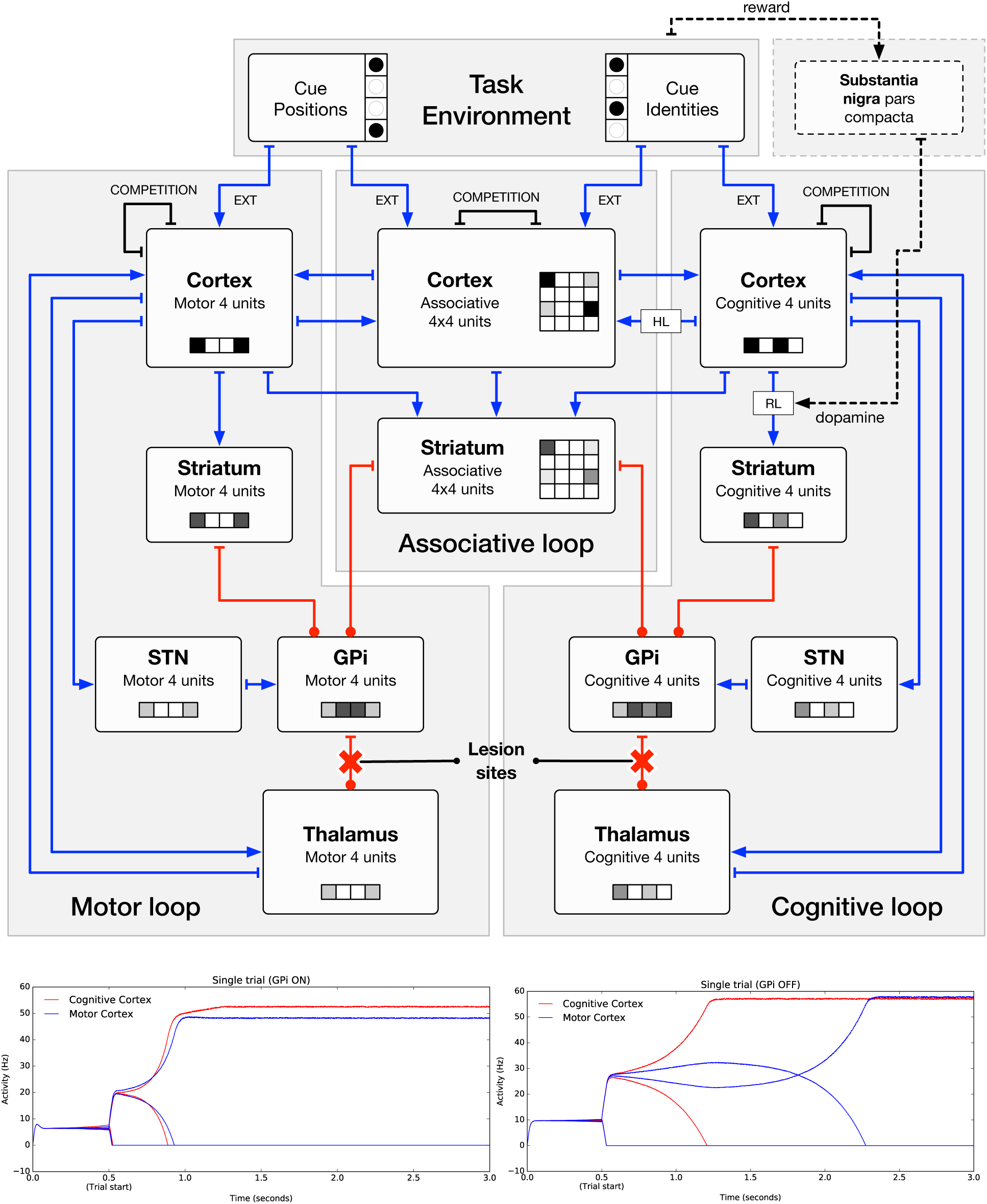
The computational model is made of 12 neural groups organized along three segregated loops (motor, associative and cognitive) that spread over the cortex, the basal ganglia and the thalamus. Blue lines represent excitatory pathways; red lines represent inhibitory pathways and dashed lines represent emulated pathways (they are not physically present in the model but their influence is taken into account). Red crosses represent lesion sites emulating the muscimol injection in the GPi of the monkeys. The color of the different units has only an illustrative purpose and does not represent actual activation. Figures from (Rougier, 2017), CC-BY.

#### Dynamics

The dynamic of a decision in the model is illustrated on the bottom part of figure 3 before any learning has occurred. The left panel shows the dynamics of the unlesioned model where a decision occurs a few milliseconds after stimulus onset. However, in the lesioned model (right panel), the suppression of the GPi output slows down considerably the decision process compared to the intact model. This means that the decision is initially driven by the basal ganglia as it has been suggested in (Pasupathy and Miller, 2005; Turner and Desmurget, 2014)

**Figure 3.**
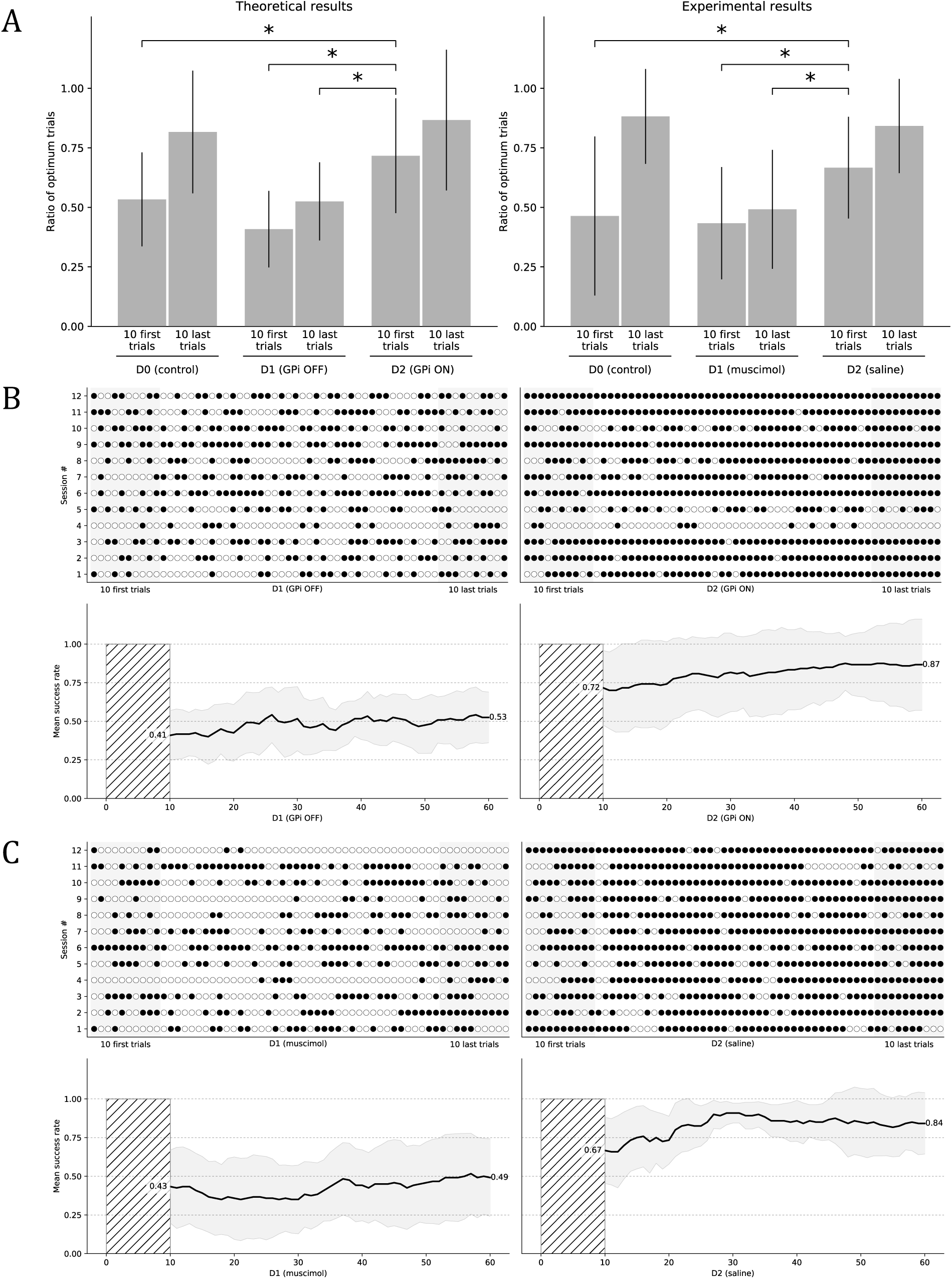
**A.** Histograms show the mean performance at the start and the end of a session in day 1 and day 2 conditions for both the model (left part) and the monkeys (right part). At the start of day 2, the performance for both the model and the monkeys is significantly higher compared to the start and end of day 1, suggesting some covert learning occurred during day 1 even though performances are random during day 1. **B & C** panels show all the individual trials (n=2x60) for all the sessions (n=12) for the theoretical model (panel **B**) and the experiments with monkeys (panel **C**). A black dot means a successful trial (the best stimulus has been chosen) and an outlined white dot means a failed trial (the best stimulus has not been chosen). Measure of success is thus independent of the actual reward received after having chosen one of the two stimuli. The bottom part shows the mean success rate over a sliding window of ten consecutive trials and averaged across all the sessions. The thick black line is the actual mean and the gray-shaded area represents the standard error of the mean (STD) over sessions. These results show that both the model and the monkeys are unable to choose the best stimulus when the GPi is disabled on day 1. This can be seen on the left part of both panels with a mean success rate oscillating around 0.5 (i.e. random choice). However, on day 2, when the GPi is re-enabled, there is an immediate effect (right part of both panels) and the mean success rate is instantly above 0.5 even if it isn’t instantly optimal. Our hypothesis is that the model benefits from the covert learning happening inside the basal ganglia during day 1 even though the BG cannot influence behavior during this period. The other consequence of this covert learning is a better estimation of both stimuli value during day 1 because there is no selection bias in favor of one or the other stimulus. They are both sampled uniformly, leading to a quantitatively uniform estimation of their respective value. * = p < 0.01. Figure from (Rougier, 2017), CC-BY.

#### Learning

Dopamine modulates learning using reinforcement learning (RL) between the cognitive cortex and the cognitive striatum such that the decision made at the cognitive level can be used to bias the decision at the motor level. Hebbian learning (HL) occurs just after a motor action has been selected and carried out and modifies the connections (LTP) between the cognitive cortex and the associative cortex. It *does not depend* on reward but only on the actual cognitive and motor choices. It is to be noted that the cortical selection (resulting from lateral competition in the cortex) is slower than the cortico-basal selection such that the cortex is initially driven by the basal ganglia output (GPi), hence it learns from the statistics provided by the BG selection.

#### Lesion

Lesion in the model is made through the removal of all the connections between the motor (resp. cognitive) GPi and the motor (resp. cognitive) thalamus (red crosses on figure 3). This prevents any communication from the basal ganglia to the model but keeps intact the communication from the cortex to the basal ganglia.

#### Statistical Analysis

Theoretical and experimental data were analyzed using Kruskal-Wallis rank sum test between the two conditions (muscimol or control) for the 6 samples (10 first trials of D0 (control), 10 last trials of D0 (control), 10 first trials of D1; 10 last trials of D1; 10 first trails of D2; 10 last trials of D2) with posthoc pairwise comparisons using Dunn’s-test for multiple comparisons of independent samples. P-values have been adjusted according to the false discovery rate (FDR) procedure of Benjamini-Hochberg. Results were obtained from raw data using the PMCMR R package (Pohlert, 2014). Significance level was set at P< 0.01.

## Results

We designed a simple two-armed bandit task where two stimuli A and B are associated with different reward probability as explained on figure 1. The goal for the subject is to choose the stimulus associated with the highest reward probability, independently of its position. The protocol is split over two days. On the first day, a novel set of stimuli (with respective reward probability 0.25 and 0.75) is used while the GPi output is suppressed and a session is made of 60 consecutive trials. On the second day, GPi suppression is removed and the same set of stimuli from day 1 is used for another 60 consecutive trials. A control has been performed using a different set of stimuli to check for the learning curve in saline condition. Note that control and two-days session uses a different set of stimuli.

**Theoretical results.** We tested our hypothesis on a computational model using 12 different sessions (corresponding to 12 different initializations of the model). The upper part of figure 3B shows individual trials for all the different sessions. On day 1, we suppressed the GPi output by cutting the connections between the GPi and the thalamus. When the GPi output is suppressed on day 1, the performance is random at the beginning as shown by the average probability of choosing the best option (expressed in mean±SD) in the first 10 trials (0.408 ±0.161) and remain so until the end of the session (0.525 ±0.164). Statistical analysis revealed no significant difference between the 10 first and the 10 last trials. On day 2, we re-established connections between the GPi and the thalamus and the model has to perform the exact same task as for day 1 using the same set of stimuli. Results shows a significant change in behavior: the model starts with an above-chance performance on the first 10 trials (0.717 ±0.241) and this change is significant as compared to the beginning of D1 and as compared to the end of D1, confirming our hypothesis that the BG have previously learned the value of stimuli even though they were unable to alter behavior.

**Experimental results.** We tested the prediction on two female macaque monkeys which have been implanted with two cannula guides into the left and right GPi (see Materials and Methods section for details). In order to inhibit the GPi, we injected bilaterally a GABA agonist (muscimol, 1μg) 15 minutes before working session (see Materials & Methods) on day 1. The two monkeys were trained for 7 and 5 sessions respectively, each session using the same set of stimuli. Results on day 1 shows that animals were unable to choose the best stimulus in such condition from the start (0.433 ±0.236) to the end (0.492 ±0.250) of the session. Statistical analysis revealed no significant difference between the 10 first and the 10 last trials on day 1. On day 2, we inject bilaterally a saline solution 15 minutes before working session and animals have to perform the exact same protocol as for day 1. Results shows a significant change in behavior: animals start with an above-chance performance on the first 10 trials (P=0.667 ±0.213, as compared to the beginning of D1, as compared to the end of D1), confirming our hypothesis that the BG have previously learned the value of stimuli.

## Discussion

These results reinforce the classical idea that the basal ganglia architecture is based on an actor critic architecture where the dopamine serves as a reinforcement signal. However, the proposed model goes beyond this classical hypothesis and propose a more general view on the role of the BG in behaviour and the entanglement with the cortex. Our results, both theoretical and experimental, suggest that the critic part of the BG extends its role beyond the basal ganglia and makes it *de facto* a central component in behavior that can evaluate any action, independently of their origin. This hypothesis is very congruent with the results introduced in (Charlesworth et al., 2012) where authors show that the anterior forebrain pathway in Bengalese finches contributes to skill learning even when it is blocked and does not participate in the behavioural performance. To understand such hypothesis, it is important to reconsider how the basal ganglia forms a series of parallel loops (motor, associative, limbic) with the cortex and the thalamus. In higher order mammals, such as primates, the overall process starts in the sensory cortex, where stimuli are encoded, and ends up preferentially in the motor cortex from where an actual action is sent to the medullar motor neurons. Accordingly, in the previous versions of our model, the cortex was considered as a single input/output excitatory layer without intrinsic dynamic properties other than the I/O function of the populations. We then added a thalamic loop which allowed positive feedback but the different channels/populations were still independent. This limited autonomy is reasonable to mimic non-mammal vertebrate 3-layers dorsal mantle (aka pallium), but it is too rudimentary for the more complex architecture of the 6-layers mammal cortex. The latter is remarkable for its organization in functional columns that are able to provide themselves positive feedback and to exert lateral inhibition on their neighbors. This architecture grants the cortex with dynamic properties that are far beyond what we were able to capture with our previous versions of the model: the balance between activation and inhibition allowed to toggle into various states that could be segregated enough to trigger different decisions. We therefore decided to add a cortical module encompassing these properties. We also grant it with the capacity to perform Hebbian learning based on the consensual hypothesis that this property is shared by most of the cortical structures. This new model captures the very essence of animal behavior in our two-armed bandit like task and provides a non-intuitive prediction about the occurring of covert learning in the striatum while the output of the BG is disrupted. It also reinforces the idea we proposed in (Piron et al., 2016) that a behavioral decision results from both the cooperation (acquisition) and the competition (expression) of two distinct but entangled actors under the evaluation of a single critic. However, if the theoretical model proposes the cortex to be a potential actor, we have no experimental evidence yet that this is the case for the monkeys. From the experiments, we conducted, we can only deduce that the monkeys are able to initiate random choices following injection of muscimol in the internal part of the Globus Pallidus and that they learn the value of the different stimuli in the meantime. If the theoretical model proposes that the motor cortex is able to make such move thanks to lateral competition, it is yet to be confirmed at the experimental level.

**Table 1.**
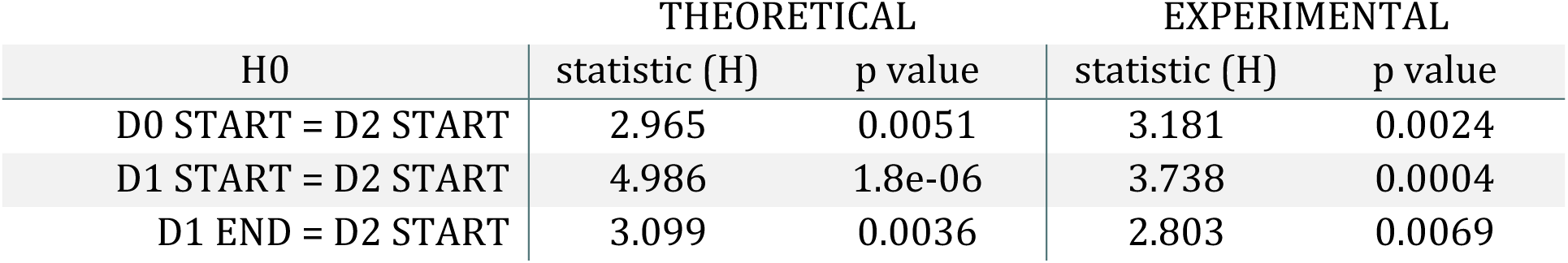
Results from the Kruskal-Wallis rank sum test with posthoc pairwise comparisons using Dunn’s-test for multiple comparisons of independent samples. P-values have been adjusted according to the false discovery rate (FDR) procedure of Benjamini-Hochberg. Results were obtained from raw data using the PMCMR R package (Pohlert, 2014).

